# Collagen Hydrogel Tube Microbioreactors for Cell and Tissue Manufacturing

**DOI:** 10.1101/2025.01.08.631570

**Authors:** Yakun Yang, Xinran Wu, Ying Pan, Yong Wang, Xiaojun Lian, Cheng Dong, Wansheng Liu, Shue Wang, Yuguo Lei

## Abstract

The production of mammalian cells in large quantities is essential for various applications. However, scaling up cell culture using existing bioreactors poses significant technical challenges and high costs. To address this, we previously developed an innovative 3D culture system, known as the AlgTube cell culture system, for high-density cell cultivation. This system involves processing cells into microscale alginate hydrogel tubes, which are suspended in the culture medium within a vessel. These hydrogel tubes shield cells from hydrodynamic stress and maintain the cell mass below 400 µm in diameter, facilitating efficient mass transport and creating a favorable microenvironment for cell growth. Under optimized conditions, AlgTubes supported long-term culture with high cell viability, rapid expansion (1000-fold increase over 9 days per passage), and high yield (5×10⁸ cells/mL), offering significant advantages over conventional methods. Despite these benefits, AlgTubes have critical drawbacks. They are mechanically fragile, with frequent breakage during culture leading to cell leakage and production failures. Additionally, many cell types exhibit poor growth due to the inability to adhere to the alginate surface, making alginate hydrogel microtubes unsuitable for industrial-scale cell production. To overcome these challenges, we developed a novel collagen hydrogel tube-based microbioreactor system in this work. This system provides enhanced robustness and adhesion, enabling scalable, cost-effective, and efficient cell production for a wide range of applications.

## A. Introduction

Mammalian cells have diverse and critical applications. Stem cells, such as human pluripotent stem cells (hPSCs) - including human embryonic stem cells (hESCs) and induced pluripotent stem cells (iPSCs) - and their progenies (i.e., cells differentiated from stem cells), can be used to treat degenerative diseases, injuries, and cancers^1^. They are also valuable for modeling diseases, screening drugs, and testing the efficacy and toxicity of drugs and chemicals. Immune cells, such as T cells and natural killer cells, can be used to treat cancers^2–6^. Mammalian cells are widely used for producing recombinant proteins and viruses in both research and industrial settings, with many of these products playing essential roles in clinical applications^7–9^.

These applications require vast numbers of high-quality cells^1^. For example, approximately 10^5^ surviving dopaminergic (DA) neurons, ∼10^9^ cardiomyocytes, or ∼10^9^ β cells are needed to treat a single patient with Parkinson’s disease (PD), myocardial infarction (MI), or Type 1 diabetes, respectively^10^. Moreover, far greater quantities of cells are required initially due to low yields during in vitro culture and poor survival rates following transplantation. For instance, only <10% of transplanted dopaminergic neurons and cardiomyocytes reportedly survive in a few days after transplantation. The demand is further amplified by the prevalence of degenerative diseases and organ failure, with over 1 million people in the United States living with PD, 1–2.5 million with Type 1 diabetes, and ∼8 million with MI. Tissue engineering also requires large cell quantities. For example, approximately 10^10^ hepatocytes or cardiomyocytes are needed to construct an artificial human liver or heart^11^. Similarly, drug discovery efforts, such as screening a million-compound library, may require 10^10^ cells per screen^10^. Advances in combinatorial chemistry, RNA research, and the study of complex signaling and transcriptional networks have led to the development of extensive libraries, necessitating large-scale cell production. Large numbers of mammalian cells, such as Chinese Hamster Ovary cells (CHO cells) and Human Embryonic Kidney 293 cells (HEK293), are also needed for producing therapeutic biologics, such as monoclonal antibodies, enzymes and viral particles^7–9^.

Efficient, scalable, and cost-effective methods for producing high-quality cells, especially for clinical purposes, remain a pressing challenge^1,10,12^. In vivo, human cells exist within intricate three-dimensional (3D) microenvironments that support critical interactions between cells and the extracellular matrix (ECM), provide robust nutrient and oxygen delivery, and minimize hydrodynamic stress^13–17^. Unfortunately, current cell culture methods often fail to replicate these conditions, resulting in inefficiencies and difficulties in scaling production.

Two-dimensional (2D) cell culture systems, such as flasks, are among the most commonly used methods for cell cultivation. Despite their popularity, these systems lack the complexity of in vivo environments. They are also resource-intensive in terms of labor, space, and reagents, making them unsuitable for large-scale applications^1,10,12^. In response, 3D suspension culture techniques, including stirred-tank bioreactors, have been developed to enable larger-scale production. While promising, these systems face challenges related to uncontrolled cell aggregation, particularly in cells with strong adhesion tendencies^16,17^. In such cultures, cells form large aggregates. It is known that the transport of essential nutrients, oxygen, and growth factors to cells at the core of aggregates larger than 400 µm become difficult, leading to slower proliferation, apoptosis, and undesired differentiation in stem cells^10,18^. Although agitation can help reduce aggregation and improve mass transport, it can also introduce shear forces that negatively impact cell survival and growth^10,19,20^. Consequently, 3D suspension cultures often exhibit high cell death, slow growth rates, and low volumetric yields. For example, hPSCs typically expand only four-fold in four days, producing yields of approximately 2.0×10⁶ cells/mL—utilizing just 0.4% of the bioreactor volume^21–23^.

The complex hydrodynamic conditions in stirred-tank bioreactors - such as flow direction, shear stress, and chemical gradients - add further complications. These variables depend on multiple factors, including the bioreactor’s design, the viscosity of the medium, and the agitation speed, making precise control difficult^1,10,12,19–24^. Studies on producing cardiomyocytes from hPSCs in stirred-tank bioreactors highlight these challenges. Yields from three ∼100 mL batches of human embryonic stem cells (hESCs) ranged from 40 to 100 million cells, with cardiomyocyte purity varying between 54% and 84%. Using a different hPSC line under the same conditions produced yields between 89 and 125 million cells, with purity varied from 28% to 88%^25,26^. Scaling up from ∼100 mL to ∼1000 mL would require extensive re-optimization of bioreactor design, agitation rates, and protocols, underscoring the technical and economic difficulties of achieving industrial-scale production. Currently, the largest demonstrated hPSC suspension cultures remain limited to volumes of tens of liters^10,27^.

We proposed to develop hydrogel tube-based 3D microbioreactors to address these limitations. Previously, we demonstrated that growing cells in hollow hydrogel microtubes or microbioreactors made from alginate polymer could address many of these challenges. These microtubes prevent excessive aggregation, ensure efficient mass transport to cells, and eliminate hydrodynamic stress. This approach supports high cell viability, rapid proliferation, and high volumetric yields (e.g., up to 5×10^8^ per milliliter of volume), dramatically reducing bioreactor volume, production time, and cost^28–37^.

Despite their advantages, alginate hydrogel microtubes have critical limitations. They are mechanically fragile, with a significant proportion breaking during cell culture, leading to cell leakage and production failures. Additionally, many cells do not grow well in alginate hydrogel microtubes because they cannot adhere to the alginate surface. Therefore, alginate hydrogel microtubes are not suitable for industrial cell production. There is a critical need for new, robust, and adhesive hydrogel tube microbioreactors. In this study, we present a novel collagen hydrogel tube-based microbioreactor system that addresses these limitations, enabling scalable, cost-effective, and high-efficiency cell production across diverse applications.

## B. Methods

### Collagen Extraction from Rat Tail Tendons

Rat tails were immersed in 70% ethanol to remove debris, and the skin was carefully stripped away using a scalpel and forceps. The exposed tendons were then collected and thoroughly washed three times with PBS. Subsequently, they were sterilized in 70% ethanol for a minimum of 1 hour. The sterilized tendons were dissolved in 0.02N acetic acid with continuous stirring at 4°C for 48 hours. The resulting viscous solution was centrifuged at 10,000 rpm for 60 minutes at 4°C. The supernatant obtained from centrifugation was dialysis against 0.02 acetic acid and collected as the collagen stock solution for further applications.

### Processing Collagen Tubes

A custom micro-extruder was designed using Fusion 360 (Autodesk) and fabricated using a stereolithography 3D printer (Form 3B+, Formlabs) for collagen tube production. The three inlets were connected to syringes mounted on syringe pumps (Fusion 200, Chemyx) to precisely control flow rates. A syringe containing a 1.5% methylcellulose (MC) or hyaluronic acid solution with single cells was connected to the core flow channel of the micro-extruder and pumped at a rate of 30 μL/min. A pre-chilled syringe loaded with collagen solution was placed in a custom ice box, connected to the shell flow channel, and pumped at a rate of 180 μL/min. A syringe containing 50 mM HEPES buffer was attached to the sheath flow channel and pumped at a rate of 2 mL/min. The outlet of the micro-extruder was submerged in a 37°C, 50 mM HEPES buffer maintained using a heating pad. Once the pumps were activated, collagen tubes were continuously generated at the outlet and collected into a tube filled with 50 mM HEPES buffer. Subsequently, the 37°C HEPES buffer was replaced with pre-heated cell culture medium to culture cells.

### Confocal Microscopy Imaging

Bulk collagen was prepared using rat-tail collagen and a neutralization solution. Briefly, collagen was neutralized by adding the neutralization solution at a 9:1 ratio. The mixture was then incubated at 37°C for 15 minutes to allow gelation. To visualize the nanostructures of bulk collagen and ColTubes, ATTO 488 NHS ester was used to label the collagen fibers, following the manufacturer’s guidelines. Briefly, a 10 mM stock solution of ATTO 488 NHS ester was diluted in PBS to a final concentration of 1 µM. Bulk collagen and ColTubes were stained for 15 minutes at room temperature, followed by two washes with PBS to remove excess dye. Confocal images were captured using an Olympus FV3000 confocal microscope equipped with a 60× objective.

### Scanning Electron Microscope (SEM) Imaging

The nanostructures of collagen samples were analyzed using a Zeiss SIGMA VP-FESEM (SEM). Prior to imaging, the samples were dehydrated through a graded ethanol series (25%, 50%, 70%, 85%, 95%, and 100%), spending 5 minutes at each concentration. This was followed by critical point drying using a Leica EM CPD300 Critical Point Dryer. To enhance conductivity, the samples were sputter-coated with a 4.5 nm layer of iridium using a Leica EM ACE600 Sputter Coater. SEM images were acquired under high vacuum conditions at an accelerating voltage of 5 kV, with a working distance of 6 mm. Magnifications ranged from ×150 to ×5,000 to visualize the collagen fiber structures in detail.

### Doping Collagen Tubes with Extracellular Matrix (ECM) Proteins

Recombinant Human Laminin 511 E8 fragments (iMatrix-511 SILK) were used as the ECM protein in this experiment. For staining the ECM protein, ATTO 594 NHS ester was prepared as a 10 mM stock solution in anhydrous DMSO. The pH of the Laminin 511 E8 fragments was adjusted to 8.0–9.0 using pH 9 phosphate buffer to facilitate the reaction with ATTO 594 NHS ester. The dye stock solution was then added to the Laminin 511 E8 fragments solution at a 100:1 molar ratio (Dye:Protein). The mixture was vortexed at room temperature for 60 minutes and stored at 4°C for at least 48 hours before use. A PBS buffer was prepared in parallel as a control. To dope collagen tubes with ECM proteins, the acid-stabilized collagen solution was combined with the Laminin 511 E8 fragments solution on ice. For the control, the PBS buffer replaced the Laminin 511 E8 fragments in the mixture. The resulting mixture was processed into collagen tubes following the previously described method.

### Culturing hPSCs in Collagen Tubes

For a typical cell culture, 20 µL of cell solution within Collagen Tubes was suspended in 2 mL of E8 medium supplemented with 10 μM Y-27632. The cultures were maintained in a 6-well plate and incubated at 37 °C with 5% CO₂ and 21% O₂. The medium was replaced daily to ensure optimal growth conditions. To passage cells, the medium was removed, and collagen tubes were dissolved by incubating with 0.2 mg/mL Collagenase P for 15 minutes. The cell mass was collected by centrifugation at 100 g for 5 minutes, followed by treatment with Accutase at 37 °C for 10 minutes. The cells were then dissociated into single cells for subsequent culture or cryopreservation.

### Staining, Flow Cytometry, and Imaging

Single cells were fixed with 4% paraformaldehyde (PFA) at room temperature for 15 minutes, followed by incubation with PBS containing 0.1% Triton X-100, 0.5% BSA, and primary antibodies at 4°C overnight. After thorough washing, secondary antibodies were added and incubated for 2 hours at room temperature. Cells were then washed three times with PBS containing 0.5% BSA before assessment using the Attune® NxT™ Acoustic Focusing Cytometer (ThermoFisher). Data analysis was conducted using FlowJo software. LIVE/DEAD® Cell Viability staining was performed according to the manufacturer’s instructions to evaluate live and dead cells. For phase-contrast and fluorescence imaging, a Zeiss Axio Observer Fluorescent Microscope was used.

### Manufacturing Seminiferous Tubules

TM3 (Leydig) and TM4 (Sertoli) cells were cultured according to the protocol provided by ATCC. Cells were maintained in DMEM/F12 supplemented with 5% horse serum and 2.5% fetal bovine serum (FBS) at 37°C, 5% CO₂, and 95% humidity, with media changes every 2–3 days. TM3 cells were labeled with CellTrace™ CFSE (CellTrace™ 488), and TM4 cells were labeled with CellTrace™ Far Red (CellTrace™ 633) following the manufacturer’s protocol. Cells were washed with PBS and incubated with their respective dyes for 30 minutes at 37°C. After staining, excess dye was removed with two PBS washes. The stained TM4 cells were seeded in the core flow of ColTubes, while the stained TM3 cells were mixed with the shell flow of ColTubes. Fluorescence images were acquired using an Olympus FV3000 confocal microscope with a 20× objective lens.

### Statistical Analysis

All the data were analyzed using GraphPad Prism 8 statistical software and shown as mean ± standard error of the mean. P value was determined by one-way analysis of variance (ANOVA) for comparison between the means of three or more groups, log-rank test for survival, or unpaired two-tailed t-tests for two groups analysis. The significance levels are indicated by p-value, *: p<0.05, **: p<0.01, ***: p<0.001.

## C. Results

### The Extrusion System for Processing Collagen Hydrogel Tubes (ColTubes)

A novel micro-extruder with three inlets and one outlet was designed specifically for processing ColTubes (**Fig. 1A**). The extruder was fabricated using a Formula 3B printer and clear resin. A cooling box was also designed and fabricated using 3D printing (**Fig. 1B**). The ColTube processing setup consists of three syringe pumps with syringes, the micro-extruder, the cooling box, a heating pad, and a conical tube or container containing HEPES buffer (**Fig. 1C**). The three syringes contain the following solutions: (1) a cell solution at room temperature (RT, pH = 7.4), (2) an ice-cold collagen solution (pH = 3.0), and (3) a HEPES buffer (RT, pH = 7.4). The cooling box has a dedicated channel for holding syringe 1 (Fig. 1B). Ice is loaded into the box to maintain the collagen solution in syringe 1 at a temperature below 4°C. The collagen is dissolved in 0.02N acetic acid to maintain a pH of 3.0, which, along with the low temperature, prevents premature collagen gelation in syringe 1.

**Figure 1.**
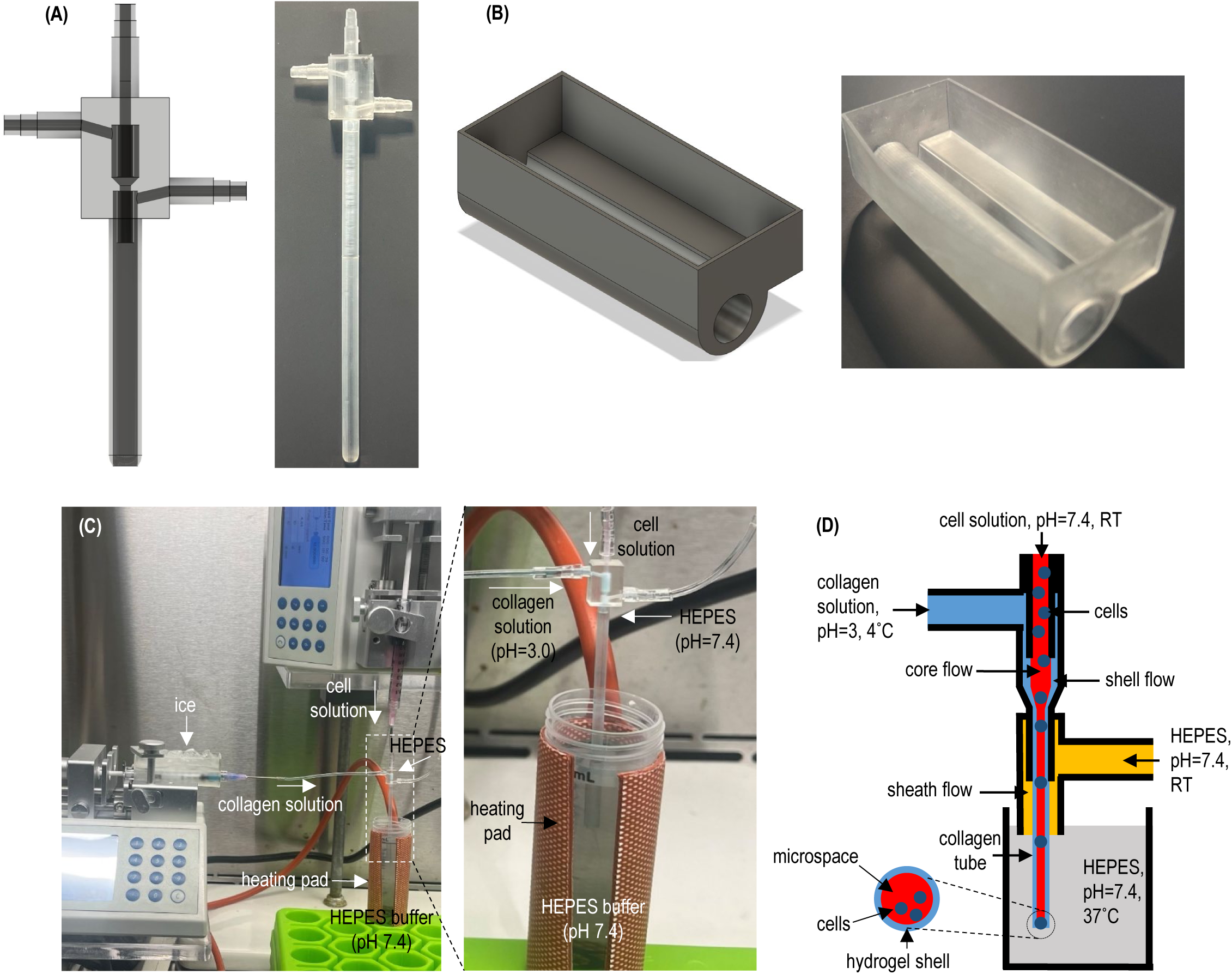
Overview of the Collagen Hydrogel Tube (ColTube) Microbioreactors. **(A)** Design and printed version of the new micro-extruder. **(B)** Design and printed version of the cooling box used for temperature control. **(C)** The setup for processing ColTubes, which includes three syringe pumps, the micro-extruder, the cooling box, a heating pad, and a HEPES buffer reservoir. **(D)** To fabricate ColTubes, a cell solution at room temperature (RT), an ice-cold collagen solution (pH = 3.0), and a HEPES buffer (RT, pH = 7.4) are pumped into the central channel and two side channels of the micro-extruder, respectively. This configuration creates coaxial core-shell-sheath flows that are extruded into a heated HEPES buffer (37°C, pH = 7.4). The shell collagen flow rapidly forms a hydrogel tube as a result of the simultaneous pH neutralization and temperature increase.

To produce ColTubes, the three solutions are pumped into inlets 1, 2, and 3 of the micro-extruder, respectively, forming coaxial core-shell-sheath flows that are extruded into the heated HEPES buffer (37°C, pH = 7.4) (**Fig. 1D**). The core, shell, and sheath flow contains the cell solution, collagen solution, and HEPES buffer, respectively. At the microscale, the extruder facilitates the formation of stable laminar flows of the three solutions. The low-pH collagen solution in the shell flow is neutralized by the core cell solution and the HEPES buffer in both the sheath flow and the conical tube. Additionally, the collagen solution is rapidly heated by the HEPES buffer in the conical tube, resulting in rapid gelation of the collagen shell, thereby forming a stable collagen tube.

In summary, our innovative micro-extruder, combined with the cooling box and heating pad setup, enables the formation of stable core-shell-sheath coaxial laminar flows while achieving rapid pH and temperature changes in the collagen flow. These innovations allow for the efficient production of microscale ColTubes suitable for cell culture. To the best of our knowledge, no other existing scalable technology can rapidly process microscale tubes with collagen without compromising cell viability.

### Engineering Principles and Control of ColTube Dimensions

ColTubes are designed to overcome the limitations of current cell culture technologies by providing physiologically relevant microenvironments for cell growth (**Fig. 2A**). In this system, cells are cultured within microscale ColTubes suspended in a culture medium inside a vessel. The hydrogel tubes create cell-friendly microspaces that promote cell-cell interactions and expansion while protecting the cells from hydrodynamic stresses within the culture vessel. The tubes also confine the cell mass to a radial diameter of less than 400 µm, ensuring efficient nutrient and waste transport throughout the culture period, as the diffusion limit in human tissue is approximately 500 µm. Additionally, the collagen tube wall is highly porous, allowing medium components to pass through freely. Finally, the collagen fibers within the tube walls serve as a substrate for cell adhesion, further supporting cell growth and functionality.

**Figure 2.**
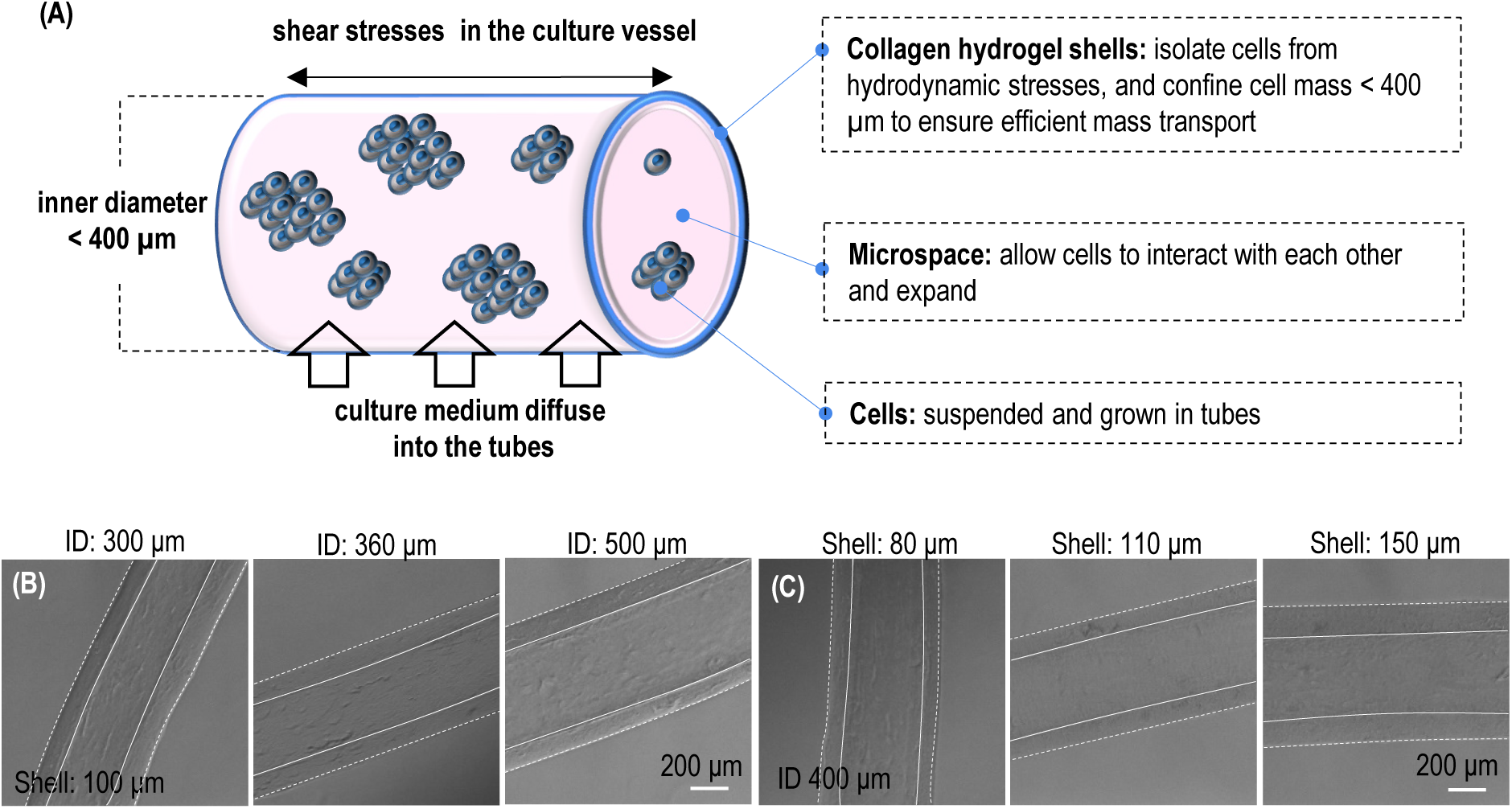
Engineering Principles and Control of ColTube Dimensions. **(A)** *Engineering Principles of ColTube Microreactors*: ColTubes are designed to culture cells within hydrogel tubes suspended in a culture medium inside a vessel. These tubes protect cells from hydrodynamic stresses and restrict the cell mass to a radial diameter of less than 400 µm, ensuring efficient nutrient and waste transport. The hydrogel tubes create cell-friendly microenvironments that facilitate cell-cell interactions and promote cell growth. Additionally, the culture medium efficiently diffuses through the porous hydrogel shell, nourishing the cells, while the collagen fibers within the tube walls provide a substrate for cell adhesion, further enhancing cell growth and functionality. **(B, C)** *Controlling ColTube Dimensions*: The dimensions of ColTubes can be adjusted by modulating the flow ratios during fabrication. In (B), the inner diameter (ID) is varied from 300 to 500 µm, while maintaining a constant shell thickness of 100 µm. In (C), the shell thickness is adjusted from 80 to 150 µm, while keeping the ID constant at 400 µm.

The dimensions of ColTubes can be precisely controlled by adjusting the flow rates of the core, shell, and sheath streams. The HEPES sheath flow not only neutralizes the acidic collagen shell solution but also acts as a hydrodynamic focusing mechanism to control the ColTube diameter. Increasing the sheath flow rate reduces the ColTube diameter, while the wall thickness can be adjusted by modifying the collagen shell flow rate. A lower shell flow rate results in a thinner tube wall. **Figures 2B** and **2C** illustrate ColTubes with varying diameters and wall thicknesses, demonstrating the versatility of the system in achieving desired tube dimensions.

### Nanostructures of ColTubes

ColTubes were successfully processed using the setup. Although the setup includes a heating pad to maintain the HEPES buffer at 37°C, ColTubes could also be processed at room temperature without the heating pad. To visualize the ColTubes, they were labeled with NHS-488 dye and imaged using a confocal microscope (**Fig. 3A**). For comparison, bulk collagen hydrogels were also prepared and imaged (**Fig. 3B**). The ColTubes exhibited collagen nanofibers forming a dense network, similar to the bulk hydrogels. The nanostructures of the outer surface and the tube wall were consistent. There are some loose collagen nanofibers protruding inward at the inner surface. Collagen concentrations in the tested range (3 - 8 mg/mL) had minimal influence on the nanofiber diameter, length, orientation, porosity, and pore size of the nanofiber network. Additionally, for ColTubes processed with a 6 mg/mL collagen concentration, there was no significant difference in nanostructures between those made at 37°C and those made at room temperature.

**Figure 3.**
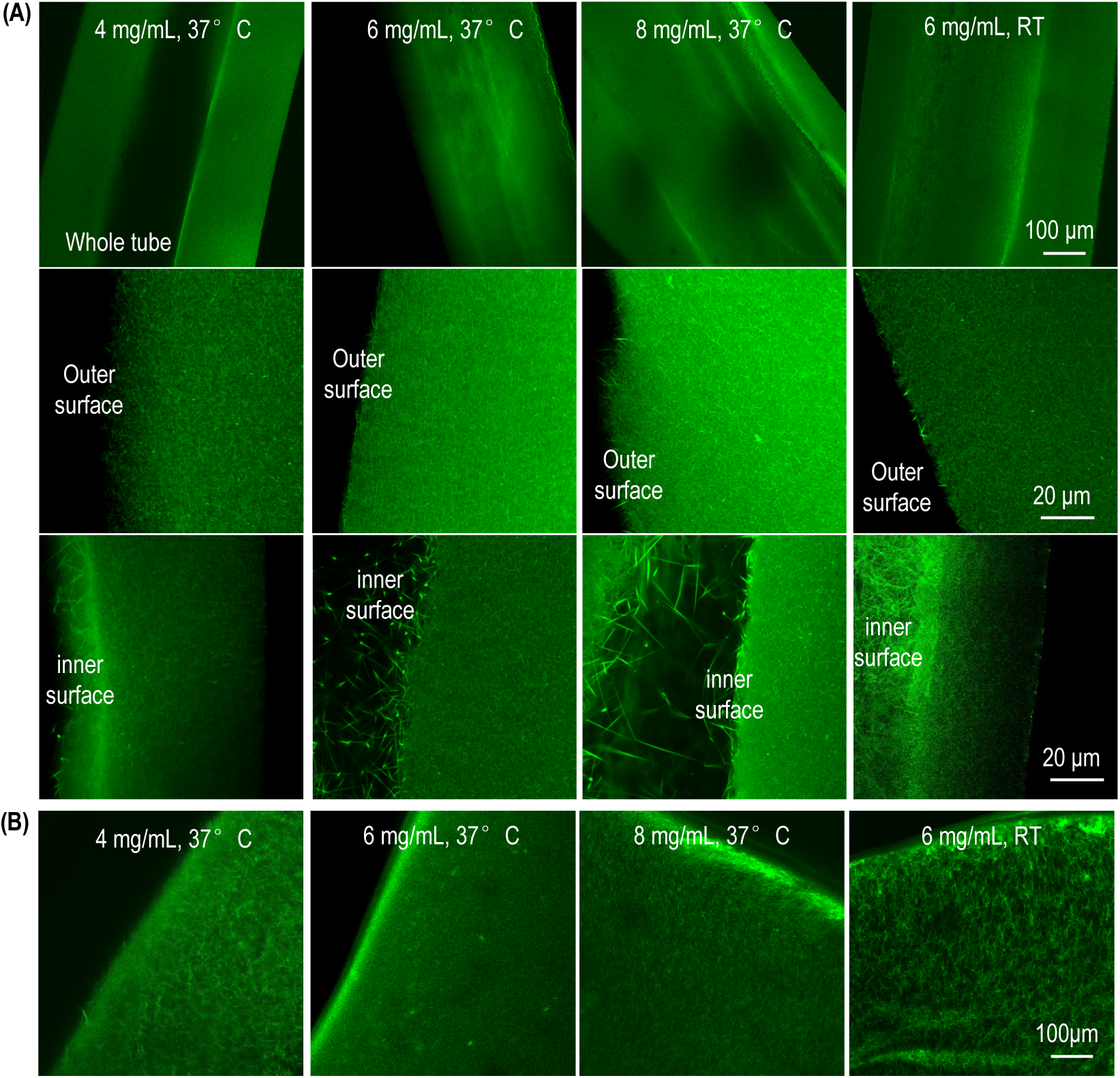
Nanostructures of ColTubes and Bulk Collagen Hydrogels. **(A)** ColTubes and **(B)** bulk collagen hydrogels were fabricated using 4-, 6-, and 8 mg/mL collagen 37°C and at room temperature. Nanostructures were imaged using confocal microscopy. For ColTubes, images include the entire tube, as well as the outer and inner surfaces.

To further investigate the structure of ColTubes, scanning electron microscopy (SEM) was performed. ColTubes were processed by fixation with paraformaldehyde (PFA), dehydration through a graded ethanol series, metal sputtering, and subsequent SEM imaging. The overall tube structure, including the inner surface, outer surface, and wall nanostructures, was visualized (**Fig. 4A**). For comparison, bulk collagen hydrogels were also prepared and imaged (**Fig. 4B**). The SEM images revealed well-defined collagen nanofibers forming a dense network within the ColTube walls. At the inner and outer surfaces, the fibers were less densely packed, with some fibers protruding inward on the inner surface, consistent with confocal microscopy observations. No notable differences in nanostructures were observed between ColTubes and bulk collagen hydrogels. Moreover, the collagen concentrations tested had minimal impact on the nanofiber diameter, length, orientation, porosity, or pore size within the nanofiber network. Additionally, ColTubes processed with a collagen concentration of 6 mg/mL exhibited no significant structural variations when fabricated at 37°C versus room temperature.

**Figure 4.**
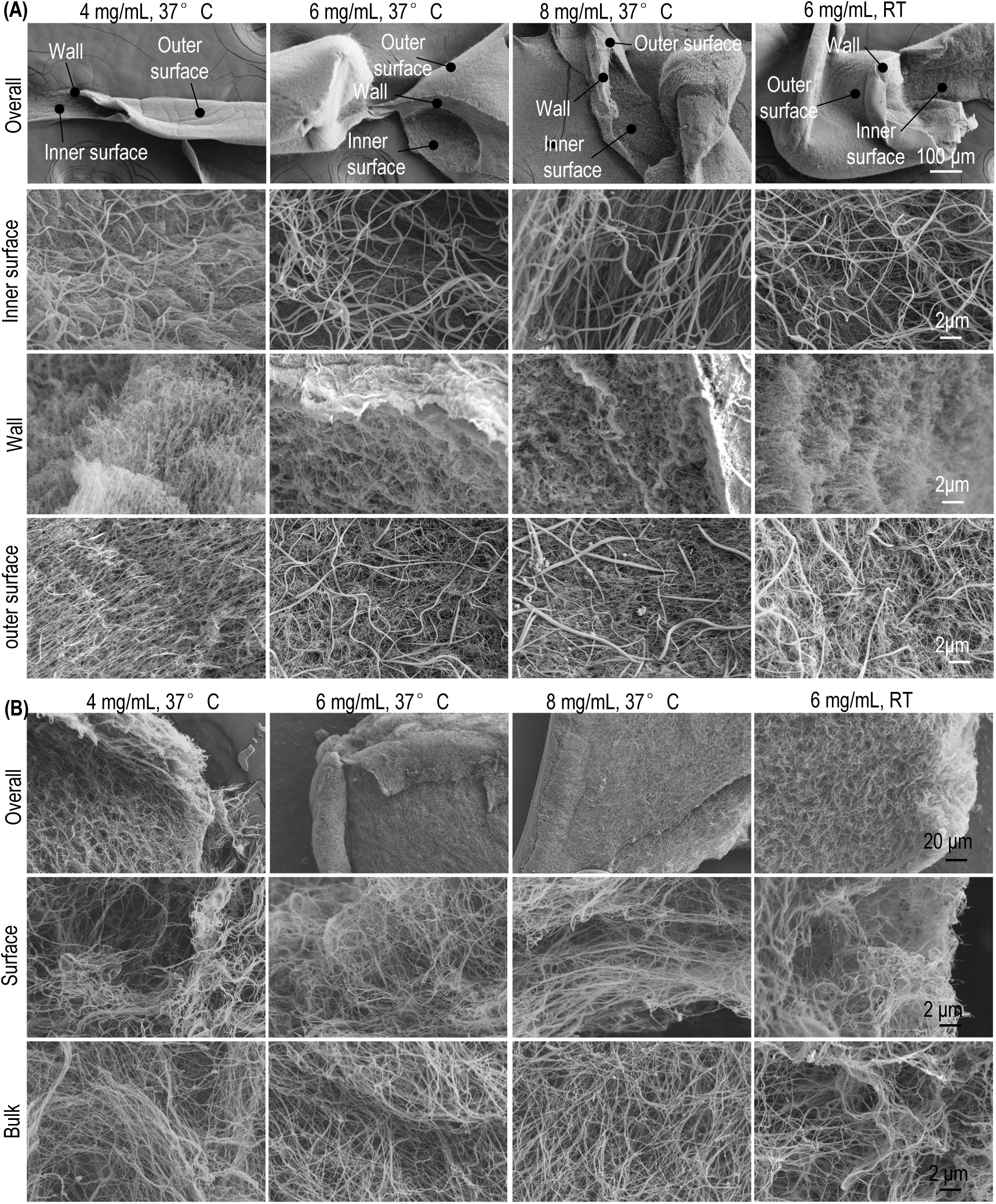
Nanostructures of ColTubes and Bulk Collagen Hydrogels. **(A)** ColTubes and **(B)** bulk collagen hydrogels were prepared using 4-, 6-, and 8 mg/mL collagen at room temperature and 37°C. Nanostructures were visualized using scanning electron microscopy (SEM). For ColTubes (A), images include the entire tube, inner surface, wall, and outer surface. For bulk hydrogels (B), both bulk and surface structures are shown.

We anticipate that cells will attach to the collagen tubes during culture. However, certain stem cells require specific ECM proteins for adhesion. For example, human pluripotent stem cells (hPSCs) are often cultured on plates coated with Laminins. To test whether ECM proteins can be incorporated and maintained in ColTubes, we labeled recombinant Laminin 511 protein with Alexa Fluor 594 NHS Ester, emitting red fluorescence. Labeled Laminin was mixed with collagen to process ColTubes, which were then stored in PBS for three days to wash away soluble Laminins. The tubes were subsequently fixed with 4% PFA and labeled with Alexa Fluor™ 488 NHS Ester to visualize both collagen and Laminin in green fluorescence. The results demonstrated that Laminin was successfully incorporated into the ColTubes and remained stable (**Fig. 5A**). Control ColTubes without Laminin did not exhibit any red fluorescence signal (**Fig. 5B**). SEM analysis revealed that doping with Laminin did not alter the nanostructures of the ColTubes (**Fig. 5C, D**).

**Figure 5.**
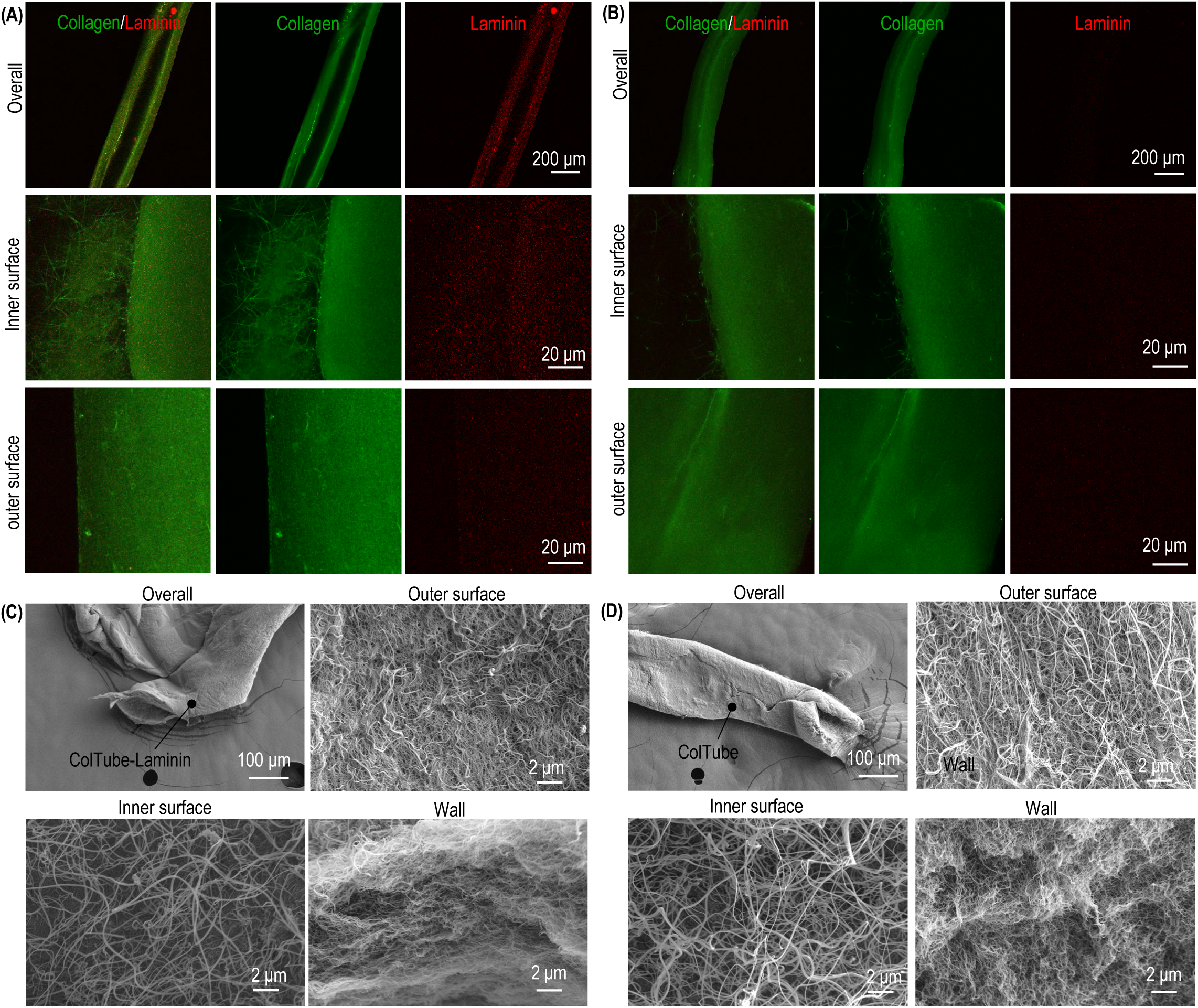
ColTubes Doped with Extracellular Matrix (ECM) Proteins. Confocal microscopy and SEM were used to analyze ColTubes with or without laminin doping: **(A)** Confocal images of the whole tube, inner surface, and outer surface of ColTubes doped with laminin. **(B)** Confocal images of ColTubes without laminin doping. **(C)** SEM images showing the whole tube, outer surface, inner surface, and wall of ColTubes doped with laminin. **(D)** SEM images of ColTubes without laminin doping.

### Culturing Cells in ColTubes

HEK293 cells are widely utilized for producing proteins and viral therapeutics. The original adherent HEK293 cells have also been adapted to suspension culture in bioreactors. We tested the suitability of ColTubes for culturing both adherent (**Fig. 6A**) and suspension (**Fig. 6B**) HEK293 cells. After 24 hours, both types of cells adhered to the inner surface of the collagen tubes and expanded to form colonies attached to the tubes. Over time, the cells proliferated further, filling the entire tube and forming a 3D cell mass. Live/Dead cell staining confirmed that the majority of cells remained viable, and we were able to harvest more than 3×10^8^ cells from one milliliter of microspaces.

**Figure 6.**
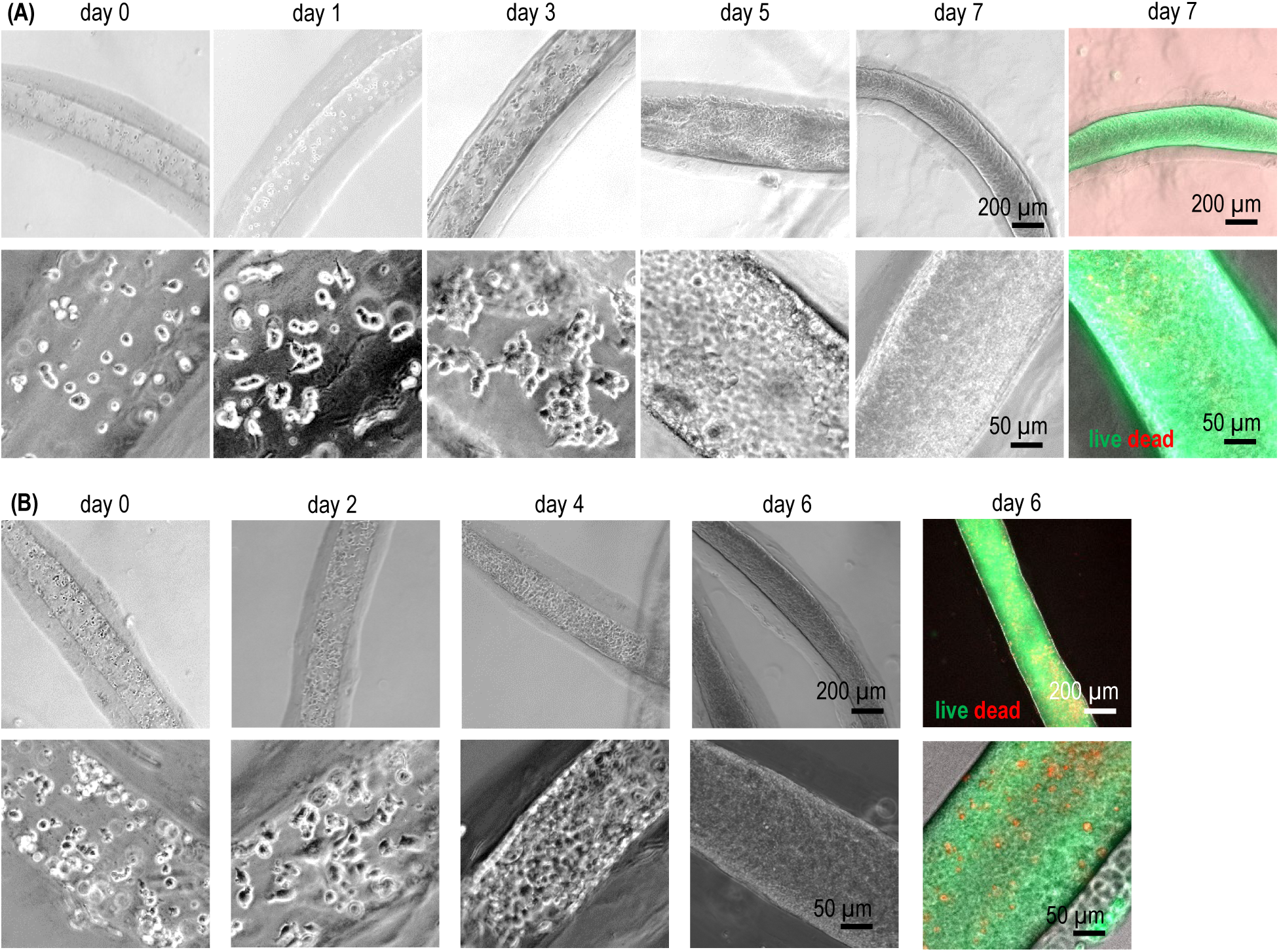
Culturing HEK 293T Cells in ColTubes. Phase-contrast and Live/Dead staining images of adherent HEK 293 cells **(A)** and suspension 293T 17SF cells **(B)** cultured in ColTubes at different time points.

Human pluripotent stem cells (hPSCs), including human embryonic stem cells (hESCs) and induced pluripotent stem cells (iPSCs), hold significant potential for treating various human diseases due to their unlimited expansion and ability to differentiate into nearly all human cell types. We cultured H9 hESCs and two iPSC lines (iPSC6 and iPSC9) in ColTubes. Single cells were processed into the tubes and attached to the inner surface within 24 hours. By day 3, cells formed small colonies attached to the tube inner surface, which grew into spheroids by day 5. By day 7, the cells had proliferated to fill most of the tubes (**Fig. 7A**). Live/Dead staining of cells before and after release from the tubes showed minimal cell death (**Fig. 7B**). Flow cytometry confirmed that 99.6% of the cells were viable (**Fig. 7C**). After dissociating the 3D cell mass into single cells, we fixed the cells with 4% PFA and stained them for the pluripotency markers Nanog and Oct4. Results showed that 97.9% and 96.1% of the cells expressed Nanog and Oct4, respectively, indicating that the hPSCs retained their pluripotency after culturing in ColTubes (**Fig. 7D**). Using ColTubes, we achieved a cell yield of over 4.5×10⁸ cells per milliliter of microspaces - more than 200 times of the yield obtained with conventional stirred tank bioreactors.

**Figure 7.**
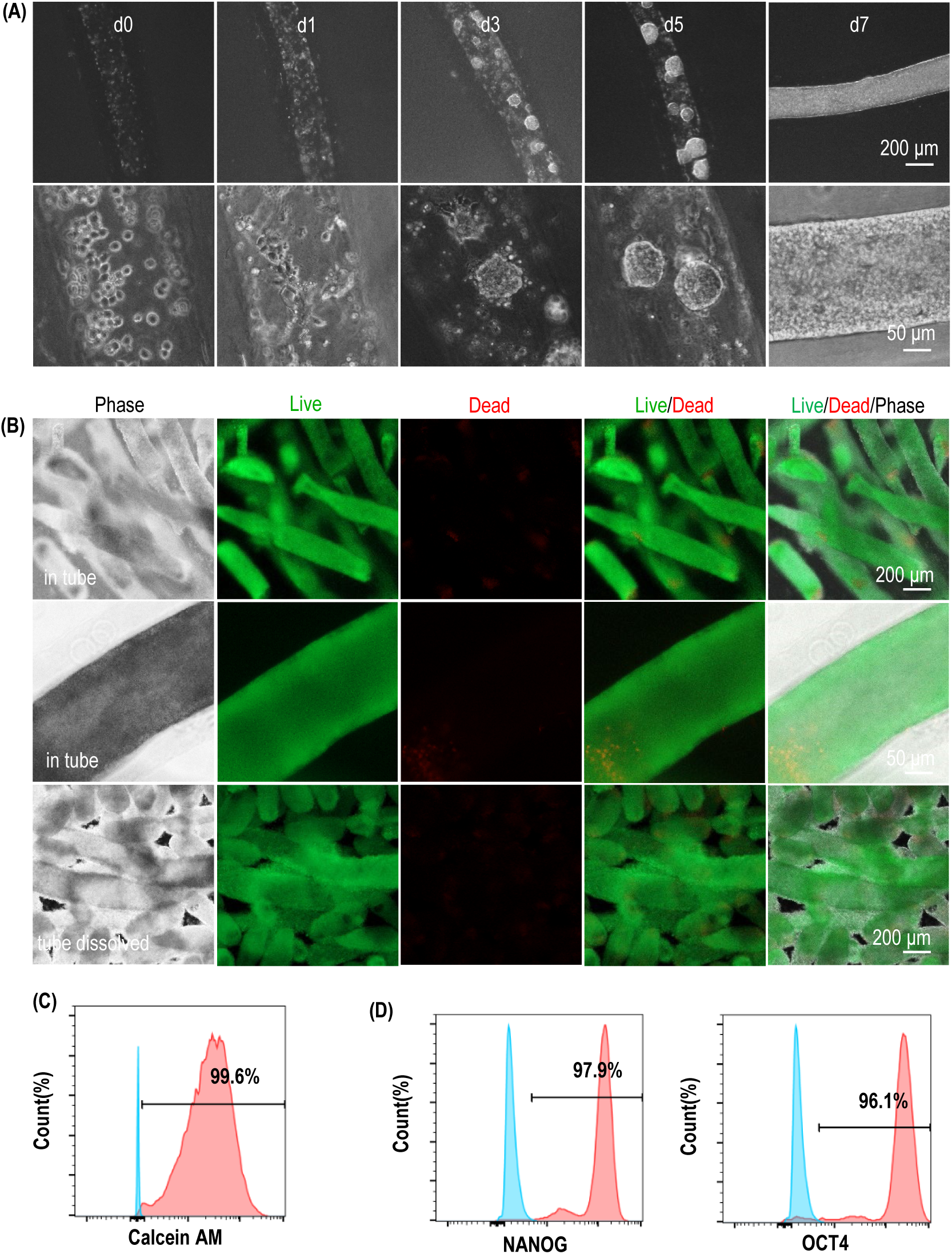
Culturing human embryonic stem cells in ColTubes. **(A)** Phase-contrast images of H9 hESCs cultured in ColTubes at days 0, 1, 3, 5, and 7. **(B)** Live/Dead staining of H9 cells on day 7 indicates high cell viability. **(C)** Flow cytometry analysis shows that 99.6% of cells harvested on day 7 are live cells. **(D)** Flow cytometry analysis confirms that the majority of cells harvested on day 7 express pluripotency markers OCT4 and Nanog.

We tested whether human pluripotent stem cells (hPSCs) could be differentiated into functional cells after expansion in ColTubes. H9 hESCs carrying a GFP reporter under the cTnT promoter were processed into ColTubes and expanded in E8 medium for 7 days. Without cell passaging, the expansion medium was switched to mesoderm induction medium to differentiate the hESCs into mesoderm progenitors. On day 2, the medium was changed to cardiac progenitor differentiation medium, followed by cardiomyocyte differentiation medium on day 7. Between days 11 and 18, cells were cultured in metabolic enrichment medium (**Fig. 8A**). Most cells remained viable, and over 98% of the final cells were cTnT+ cardiomyocytes. After treating the tubes with collagenase, fibrous cardiac tissues were harvested (**Fig. 8B**). Further dissociation of these tissues with Accutase yielded over 3×10^8^ cells per milliliter of hydrogel tubes. Therefore, ColTubes can be used for producing both microtissues and cells at high density.

**Figure 8.**
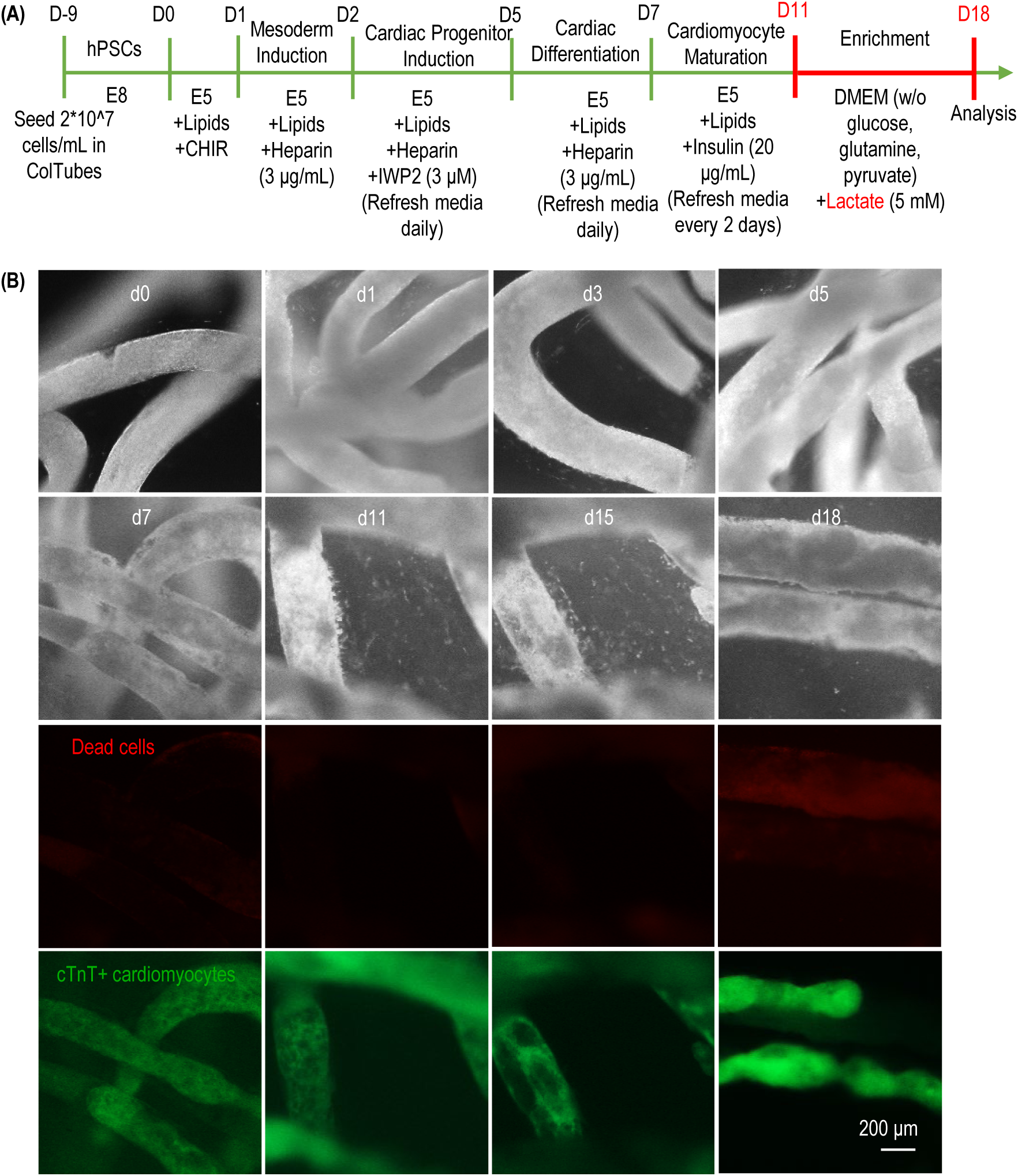
Differentiation of H9 hESCs into Cardiomyocytes in ColTubes. **(A)** Schematic illustration of the cardiomyocyte production protocol. H9 hESCs are processed into ColTubes and expanded in E8 medium, followed by mesoderm induction for 1 day, cardiomyocyte differentiation from days 2 to 11, and metabolic enrichment from days 11 to 18. **(B)** Phase-contrast and fluorescent images of cells in ColTubes on days 0, 1, 3, 5, 7, 11, 15, and 18. Cardiomyocytes are cTnT-positive, while non-cardiomyocytes are eliminated by day 18 following metabolic enrichment.

### ColTubes Exhibit Fewer Cell Leakage Events Compared to AlgTubes

Previously, we used alginate-based hydrogel tubes (AlgTubes) for cell culture. However, frequent tube breakages occurred in AlgTubes, leading to cell leakage and culture failure. As shown in **Fig. 9B**, significant cell leakage into the culture medium was observed on days 5, 6, and 7. The leaked cells formed large aggregates. In contrast, no cell leakage was observed in ColTubes (**Fig. 9A**). Quantification of leakage events (**Fig. 9C**) and the percentage of cells leaked into the medium on day 7 (**Fig. 9D**) confirmed that ColTubes exhibited no leakage events, whereas AlgTubes showed substantial leakage.

**Figure 9.**
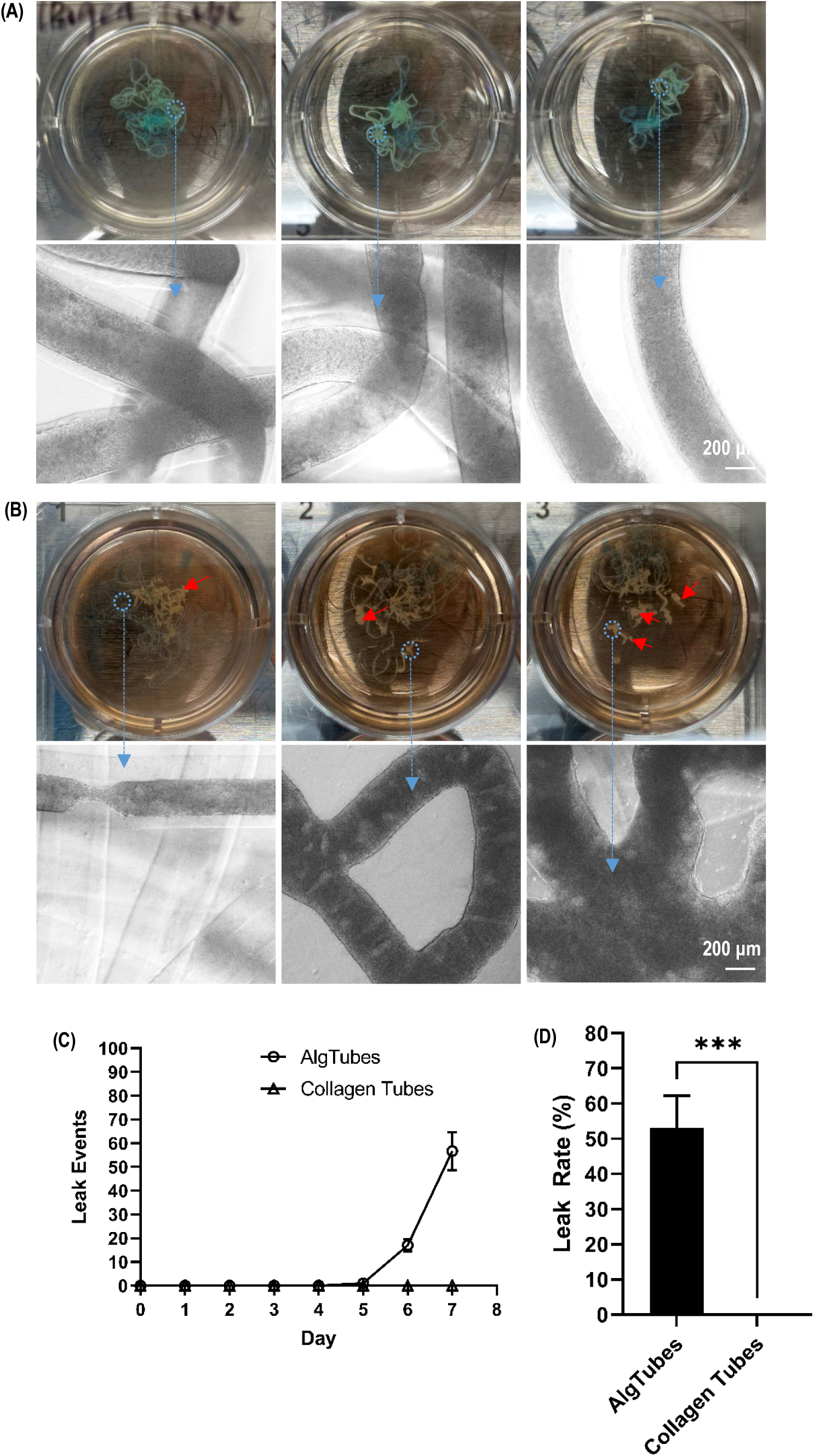
ColTubes Exhibit Fewer Cell Leakage Events Compared to AlgTubes. **(A)** Images showing no cell leakage from ColTubes. **(B)** Images showing significant cell leakage from AlgTubes, with large cell aggregates forming (red arrow). **(D)** Quantification of cell leakage events over time for AlgTubes and ColTubes. **(E)** Leak rate comparison between AlgTubes and ColTubes. Leak rate is defined as the number of cells leaked into the medium divided by the total number of cells in the well.

### Building Tubular Tissue Models with ColTubes

In addition to cell production, ColTubes can be utilized for fabricating tissues for therapeutic applications and disease modeling. Human sperm is produced in seminiferous tubules within the testis, where Sertoli cells form a monolayer lining the inner surface of the tubules, and Leydig cells, along with other minor cell types, reside in the surrounding interstitial tissue. To mimic this structure, we mixed Leydig cells with collagen to form the shell flow and suspended Sertoli cells in the core flow. The resulting ColTubes successfully contained Leydig cells within the tube walls and Sertoli cells on the inner surface, closely resembling the structure of in vivo seminiferous tubules (**Fig. 10**). Using similar methods, ColTubes can be applied to engineer other tubular tissues for various biomedical applications.

**Figure 10.**
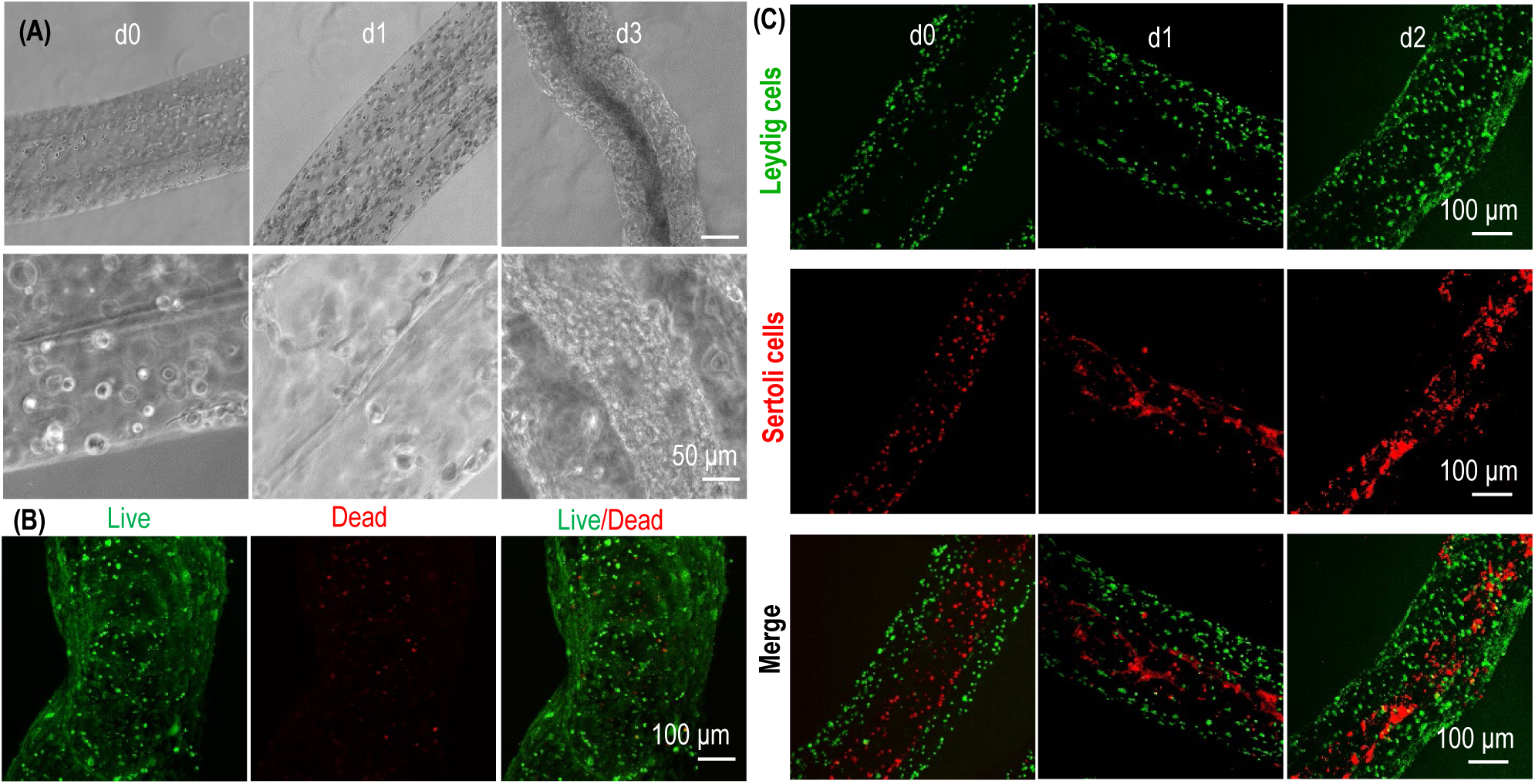
Building Seminiferous Tubules with ColTubes. **(A)** Phase-contrast images of ColTubes containing Leydig cells in the collagen wall and Sertoli cells inside the tubes at days 0, 1, and 3. **(B)** Live/Dead staining of the tubules on day 5 shows minimal cell death. **(C)** Confocal images showing Leydig cells (labeled with green dye) in the collagen wall and Sertoli cells (labeled with red dye) inside the tubes.

In summary, our results demonstrate that ColTubes are a versatile platform for both cell expansion and differentiation, as well as the production of fibrous microtissues and tubular tissues. Importantly, ColTubes exhibit excellent mechanical properties, preventing cell leakage and ensuring culture integrity. Moreover, the collagen-based structure allows cells to adhere, making ColTubes suitable for culturing both adherent and suspension cells.

## D. Discussion

Current 2D and 3D cell culture methods face significant challenges in achieving robust and cost-effective large-scale cell production^38–40^. Key issues include low cell yields, limited scalability, high costs, significant culture variability, and notable genetic and phenotypic alterations. The fundamental problem lies in the fact that the cellular microenvironments provided by these methods, such as 2D flasks and 3D stirred-tank bioreactors, differ substantially from the natural 3D microenvironments in vivo. We hypothesize that developing advanced cell culture technologies that replicate physiologically relevant microenvironments can address these limitations and substantially improve cell culture efficiency^28,31^.

We proposed that hydrogel tube microbioreactors offer a promising solution for creating cell-friendly microenvironments suitable for large-scale cell culture^28,31^. First, their hydrogel shells shield the cells inside from hydrodynamic stress, preventing stress-induced negative effects on cell survival and growth (Fig. 2A). Second, the diameter of these tubes can be precisely controlled to remain below 400 µm, ensuring that even when the tubes are densely filled with cells, the cell mass stays within the diffusion limit of human tissue. This allows efficient transport of nutrients and waste throughout the culture. Third, the hydrogel’s nanoscale pores facilitate the free passage of nutrients and growth factors from the medium through the tube walls to nourish the cells. Finally, the tubes provide a spacious environment for cells to interact and grow, which is critical for maintaining high viability, robust growth rates, and exceptional volumetric yields, as demonstrated in our previous studies.

To evaluate this approach, we developed a method to fabricate hydrogel tubes using alginate polymerst^28–37^. Alginates are widely available, inexpensive, biocompatible, and have been safely used in clinical applications. These polymers can be rapidly crosslinked with calcium ions to produce large-scale alginate hydrogel tubes (AlgTubes) through an extrusion process. After culture, AlgTubes can be dissolved with an EDTA solution that is cell-compatible, allowing for easy cell recovery. The tubes are also transparent, facilitating real-time observation of cell growtht^28–37^.

Using AlgTubes, we successfully cultured human pluripotent stem cells (hPSCs) over multiple passages with high consistency while maintaining pluripotency and chromosomal stability^28^. The cultures showed excellent viability, rapid expansion (e.g., 1000-fold increase over 10 days per passage), and exceptional volumetric yields (∼5×10⁸ cells/mL of microspace), surpassing stirred-tank bioreactors by approximately 250-fold. Expansion rates of up to 4200-fold per passage were achieved, far exceeding conventional 3D suspension cultures, which typically yield only a four-fold increase over four days. Additionally, hPSCs were differentiated into various cell types, including endothelial cells^29,32^, vascular smooth muscle cells^35^, neural stem cells^30^ and neurons^33^, all achieving yields of 5×10⁸ cells/mL. Moreover, adult cell types such as T cells were also successfully cultured in AlgTubes^34^. These results highlight the transformative potential of AlgTubes for high-efficiency, large-scale cell production.

Despite their success, AlgTubes have critical limitations. Mammalian cells lack surface receptors to bind alginate, preventing adherence to the tube walls. Many cell types, such as mesenchymal stem cells, require attachment to a substrate for survival and proliferation, making AlgTubes unsuitable for anchorage-dependent cells. Additionally, AlgTubes are mechanically fragile, leading to frequent breakages that result in cell leakage and culture failures (Fig. 9B). Further complications arise from the presence of chelators in some cell culture media, which can extract calcium ions from the alginate hydrogel and destabilize the tubes. These limitations underscore the need for further innovation in hydrogel tube design to create more robust and versatile microbioreactor systems.

In this study, we demonstrated that hydrogel tube microbioreactors made from collagen proteins could effectively address these challenges. Current technologies cannot quickly process collagen proteins into stable microtubes. Previous research has shown that collagen solutions at pH 3.0 can form hydrogels when extruded into a buffer at pH 7.4. However, fabricating stable microtubes is significantly more complex than producing solid hydrogels and had not been achieved previously. By integrating a cooling box, a three-flow micro-extruder, a heating pad, and a buffer exchange system (Fig. 1), we successfully developed a rapid method for processing collagen-based microtubes, which we call ColTubes.

The resulting ColTubes formed a dense nanofiber network that was stable in most cell culture media. They exhibited remarkable durability, with no breakages occurring during cell culture, thereby preventing cell leakage and ensuring consistent production (Fig. 9A). Furthermore, collagen’s compatibility with cellular biology provided additional advantages. Most cells express receptors for collagen, allowing them to adhere naturally (Fig. 6, 7). Additionally, collagen contains binding domains for other extracellular matrix (ECM) proteins, such as fibronectin and laminin. These proteins can be incorporated into ColTubes to support the growth of specialized cell types (Fig. 5). Our results demonstrated that cells grew in ColTubes as effectively as they did in AlgTubes.

Beyond cell culture, we also showed that ColTubes could be used to construct tubular tissue models. The human body is rich in microscale tubular tissues, such as blood capillaries, lymphatic vessels, seminiferous tubules, and milk ducts. Many cancers, including various carcinomas, arise from epithelial transformations in tubular structures, which are critical to understanding cancer development. In these tissues, endothelial or epithelial cells line the inner surfaces of the tubes, while stromal cells form the walls and the interstitial regions between tubes. ColTubes offer a platform for creating analogous tubular tissue models by placing endothelial or epithelial cells on the inner surfaces and stromal or interstitial cells within the tube walls (Fig. 10). These engineered tissues have broad applications, including disease modeling, drug screening, and regenerative medicine. They could also be used for repairing damaged tissues, providing a versatile and powerful tool for both research and therapeutic purposes.

ColTubes microbioreactors hold immense potential for widespread adoption by academic laboratories, research institutions, and biotechnology companies focused on biotherapeutics. First, ColTubes provide an efficient solution for laboratories needing to culture sufficient cells for research purposes. Most labs require fewer than 10¹⁰ cells, which can be produced using less than 20 mL of ColTubes. Additionally, ColTubes enable the culture of fibrous solid microtissues and tubular tissues, offering valuable models that mimic in vivo conditions for studying various biological processes.

Second, ColTubes have the potential to accelerate translational research. For instance, consider the development of hPSC-derived cardiomyocytes for myocardial infarction treatment^41^. Initial stages of this research typically rely on 2D culture systems to optimize differentiation protocols and establish efficacy and safety in rodent models^42^. Scaling up for preclinical large-animal studies often requires transitioning to 3D suspension cultures^43,44^, followed by further adjustments to develop large-scale bioprocesses for clinical trials^41^. Each shift in culture method necessitates extensive characterization of cell phenotype, safety, and efficacy due to bioprocess-related variations, significantly increasing the cost and timeline^45^. ColTubes offer a scalable platform suitable for all stages of this development pipeline. For example, ∼200 µL of ColTubes can suffice for protocol optimization, ∼2 mL can produce ∼10⁹ cells for small-animal studies, and ∼200 mL can yield ∼10¹¹ cells for large-animal studies and clinical trials. By eliminating the need for multiple bioprocess transitions, ColTubes can significantly reduce the time, effort, and costs involved in developing cell-based therapeutics.

Third, ColTubes present a game-changing opportunity for industrial-scale cell production. For example, producing ∼1.5×10¹⁴ hPSCs (from an initial ∼10⁸ hPSC seeding) using conventional stirred-tank bioreactors requires approximately 104,811 liters of culture volume, 11 passages, and 48 days—a technically and economically daunting task. The associated challenges include building and maintaining large bioreactor systems, extensive labor and reagent demands, and significant risks of contamination and cell loss during frequent passaging. In contrast, previous calculations using AlgTubes demonstrated that the same production could be achieved with just 320 liters of culture volume, requiring a single passage and 20 days^28^. Given that ColTubes exhibit comparable growth rates, viability, and volumetric yields to AlgTubes, they could achieve the same production scale under similar conditions. This dramatic reduction in culture volume, time, and passaging not only makes large-scale production more practical but also leads to substantial savings in labor, reagents, equipment, cGMP facility space, and overall manufacturing costs. By addressing the challenges of scalability, consistency, and cost, ColTubes have the potential to revolutionize cell culture across research, translational, and industrial applications.

In conclusion, ColTubes offer a unique combination of physiologically relevant culture microenvironments, exceptional performance and scalability, making them a promising solution for addressing the challenges of large-scale cell manufacturing. Recent advancements in biology have provided efficient protocols for differentiating hPSCs into various human cell types^1,46^. Future research should focus on incorporating these protocols into ColTubes to enable the production of diverse human cell types. Additionally, systematically investigating the potential of ColTubes for culturing other cell types, such as human adult stem cells and T cells, will further expand their utility.

## Author Contributions

Y.L. conceived the idea and directed the work. Y.L., S.W., Y.Y., X.W. designed the experiments. Y.Y., X.W., Y.P. performed experiments. Y.L., S.W., Y.Y., X.W., Y.P., Y.W., X.L., W.L., C.D. analyzed the data and wrote the manuscript.

